# Multi-omics integration of the phenome, transcriptome and genome highlights genes and pathways relevant to essential tremor

**DOI:** 10.1101/580753

**Authors:** Calwing Liao, Faezeh Sarayloo, Daniel Rochefort, Gabrielle Houle, Fulya Akçimen, Qin He, Alexandre D. Laporte, Dan Spiegelman, Alex Rajput, Patrick A. Dion, Guy A. Rouleau

## Abstract

The genetic factors predisposing to essential tremor (ET), of one of the most common movement disorders, remains largely unknown. While current studies have examined the contribution of both common and rare genetic variants, very few have investigated the ET transcriptome. To understand pathways and genes relevant to ET, we used an RNA sequencing approach to interrogate the transcriptome of two cerebellar regions, the dentate nucleus and cerebellar cortex, in 16 cases and 16 age- and sex-matched controls. Additionally, a phenome-wide association study (pheWAS) of the dysregulated genes was conducted, and a genome-wide gene association study (GWGAS) was done to identify pathways overlapping with the transcriptomic data. We identified several novel dysregulated genes including *CACNA1A*, a calcium voltage-gated channel implicated in ataxia. Furthermore, several pathways including axon guidance, olfactory loss, and calcium channel activity were significantly enriched. A subsequent examination of the ET GWGAS data (N=7,154) also flagged genes involved in calcium ion-regulated exocytosis of neurotransmitters to be significantly enriched. Interestingly, the pheWAS identified that the dysregulated gene, *SHF*, is associated with a blood pressure medication (P=9.3E-08), which is commonly used to reduce tremor in ET patients. Lastly, it is also notable that the dentate nucleus and cerebellar cortex have different transcriptomes, suggesting that different regions of the cerebellum have spatially different transcriptomes.

## Introduction

Essential tremor (ET), one of the most common movement disorders, involves rhythmic shaking during voluntary movements, particularly in the hands^1^. Although the disease is not fatal, it can have large negative effects on daily life and psychological well-being. Familial clustering suggests that genetic factors have an important role in ET. Twin studies have shown that ET has a concordance of 69–93% in monozygotic twins and 27–29% in dizygotic twins, suggesting that both genetics and environmental factors drive the phenotype^2^. Despite dozens of studies investigating the genetic etiology of ET, the heritability has largely remained unexplained. This is likely due to the misdiagnosis of ET as other similar movement disorders (e.g. Parkinson’s disease and dystonia), phenocopies, genetic heterogeneity and incomplete penetrance of risk alleles, greatly reducing statistical power of linkage studies^3,4^.

By comparison to other neurological conditions, there have been relatively few genetic studies of ET. These studies have used approaches that range from screening of function-based candidate genes, linkage and gene associations, and high-throughput sequencing of familial cases. In 2016, the largest genome-wide association study (GWAS) thus far reported used ET cases of European descent and identified three genomic loci associated with ET^5^. Since this last study, replications were undertaken across cohorts of ET cases of different ethnic origin (e.g. Han-Chinese), yet only a few successfully replicated the association of a single locus. The failure to replicate more than a locus is possibly due to relatively small cohorts and haplotype structures that were too different from the one originally used in 2016. A form of study that is absent from the ET literature is a high-throughput transcriptomic-wide approach to identify gene dysregulations across the expression profile of disease relevant brain tissue. The cerebellum has been previously implicated in ET through histological studies. Specifically, atrophy and dysregulation of Purkinje cells have largely been associated with ET^6^. Since little is known about the underlying biology of ET and genomic studies have not found adequate evidence for the suggested heritability of ET, we conducted RNA sequencing to identify dysregulated genes and pathways. We sequenced two distinct regions of the cerebellum: the cerebellar cortex and the dentate nucleus, in 16 cases and 16 age- and sex-matched controls. Additionally, we conducted a genome-wide gene association study (GWGAS) using ET GWAS data to narrow down relevant pathways. Several interesting genes such as *PRKG1* and *CACNA1A* were differentially expressed. A phenome-wide association study (pheWAS) of the differentially expressed genes (DEGs) identified blood pressure medication as a relevant phenotype for the DEG, *SHF*. Both RNA sequencing and GWGAS identified calcium-channel relevant pathways suggesting that dysregulation in these pathways contribute towards the ET phenotype.

## Results

### Differentially expressed genes identify potentially disease relevant pathways

Six genes were differentially expressed in the cerebellar cortex and two genes were differentially expressed for the dentate nucleus for an FDR of 0.05 (Table 1). The QQ-plot did not show any large stratification of the data in cerebellar cortex or dentate nucleus (Supplementary Figure 1, 2). Interestingly, the top dentate nucleus differentially expressed genes differed compared to the cerebellar cortex. The top non-coding RNA differentially expressed was *LINC00599* in the cerebellar cortex, which was highly associated with calcium ion-regulated exocytosis of neurotransmitter (P = 3.3E-06), ionotropic glutamate receptor signaling pathway (P = 9.2E-10), neurotransmitter secretion (P = 1.6E-09) and synaptic vesicle exocytosis (P=1.0E-07) in the GO database based on GeneNetwork’s co-expression database. Pathway analyses of the DEGs based on co-expression identified six clusters for the differentially expressed genes (Figure 1). Several novel pathways possibly implicated in ET were identified including axon guidance, olfactory receptor activity, and voltage-gated calcium channel activity for the cerebellar cortex (Table 2). Two gene clusters were identified for the dentate nucleus (Figure 2). Many pathways were enriched in the dentate nucleus, including olfactory transduction, olfactory signaling pathway and MAPK signaling (Table 3). Splicing analysis with rMATs and SUPPA2 did not find any significant retained introns, mutually exclusive exons, alternative 3’ splice sites, alternative 5’ splice sites or skipped exons between cases and controls. Reverse transcriptase qPCR (RT-qPCR) of the DEGs supported the direction as the Wald test statistic (Supplementary Table 1).

**Table 1.**
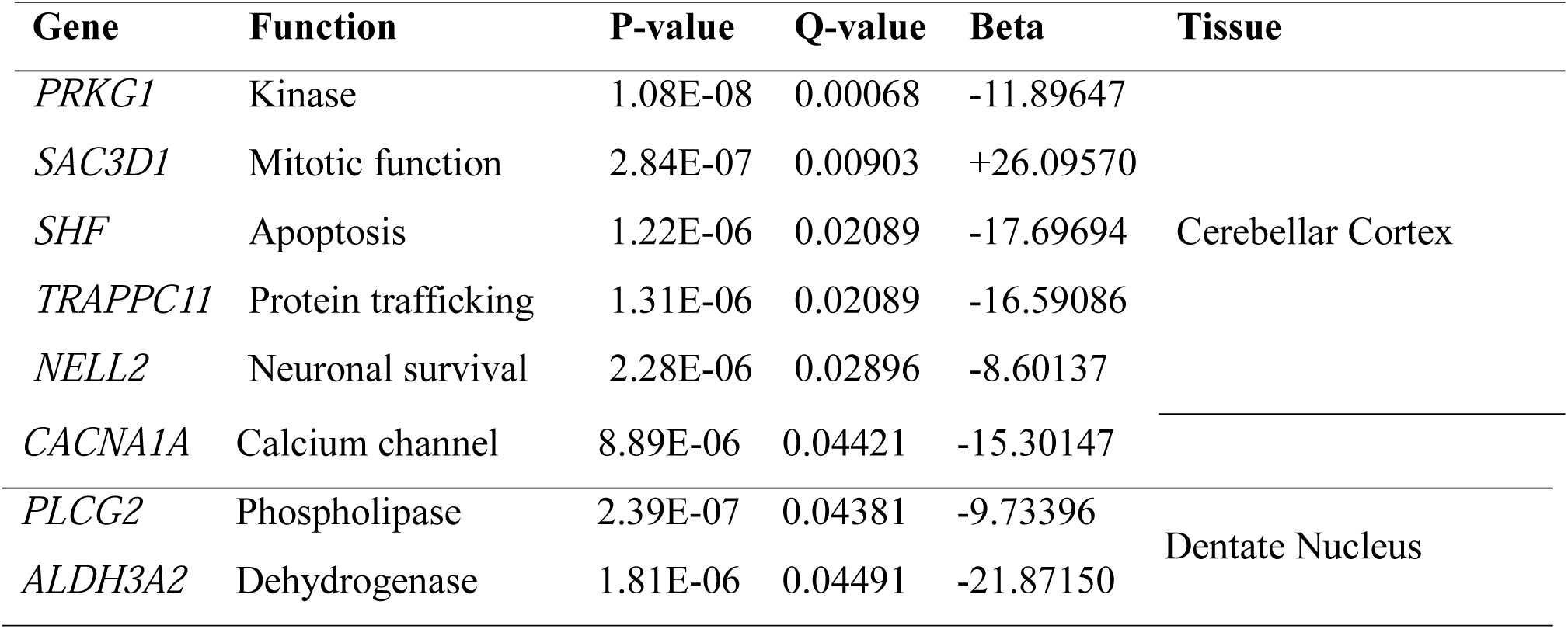
Differentially expressed genes of the cerebellum in ET patients compared to controls.

**Table 2.** Significant pathways of gene clusters identified through gene network analysis for the cerebellar cortex.

| Pathway | P-value | Database | Cluster |
| --- | --- | --- | --- |
| Cardiac conduction | 6.4E-04 | GO Processes | 1 |
| Triglyceride lipase activity | 6.9E-03 | GO Function |  |
| Olfactory receptor activity | 2.8E-03 | GO Function |  |
| Voltage-gated calcium channel complex | 3.2E-04 | GO Cellular |  |
| Positive regulation of cytosolic calcium ion concentration involved in phospholipase C-activating G-protein coupled signaling pathway | 6.5E-03 | Go Processes | 2 |
| Neuromuscular junction | 1.7E-03 | Go Cellular |  |
| Axon | 4.5E-03 | Go Cellular |  |
| Presynaptic depolarization and calcium channel opening | 2.5E-03 | Reactome | 3 |
| Transmission across chemical synapses | 1.5E-03 | Reactome |  |
| Amyloid-beta binding | 5.0E-03 | GO Function |  |
| GABA-A receptor complex | 5.7E-04 | GO Cellular |  |
| Neuron projection | 4.9E-04 | GO Cellular |  |
| Signaling by GPCR | 1.6E-04 | Reactome | 4 |
| Positive regulation of cytosolic calcium ion concentration | 4.5E-03 | Go Processes |  |
| Potassium ion leak channel activity | 5.6E-03 | GO Function | 5 |
| Vascular smooth muscle contraction | 1.6E-03 | KEGG |  |
| Transmission across chemical synapses | 4.9E-04 | Reactome | 6 |
| Presynaptic depolarization and calcium channel opening | 1.0E-03 | Reactome |  |
| Axon guidance | 3.5E-03 | Reactome |  |
| Regulation of glutamatergic synaptic transmission | 3.6E-04 | Go Processes |  |
| Cellular calcium ion homeostasis | 9.5E-03 | GO Processes |  |
| High voltage-gated calcium channel activity | 5.0E-04 | GO Function |  |
| Hypertrophic cardiomyopathy | 4.7E-04 | KEGG |  |

**Table 3.** Significant pathways of clusters with DEGs identified through gene network analysis for dentate nucleus.

| Pathway | P-value | Database | Cluster |
| --- | --- | --- | --- |
| Nuclear events mediated by MAPK | 4.6E-06 | Reactome | 1 |
| Regulation of synaptic plasticity | 6.2E-03 | GO Processes |  |
| Olfactory transduction | 7.3E-03 | KEGG |  |
| Olfactory signaling pathway | 4.9E-04 | Reactome | 2 |
| Neuron migration | 4.0E-03 | Go Processes |  |
| Protein kinase activity | 2.8E-03 | GO Function |  |

**Figure 1.**
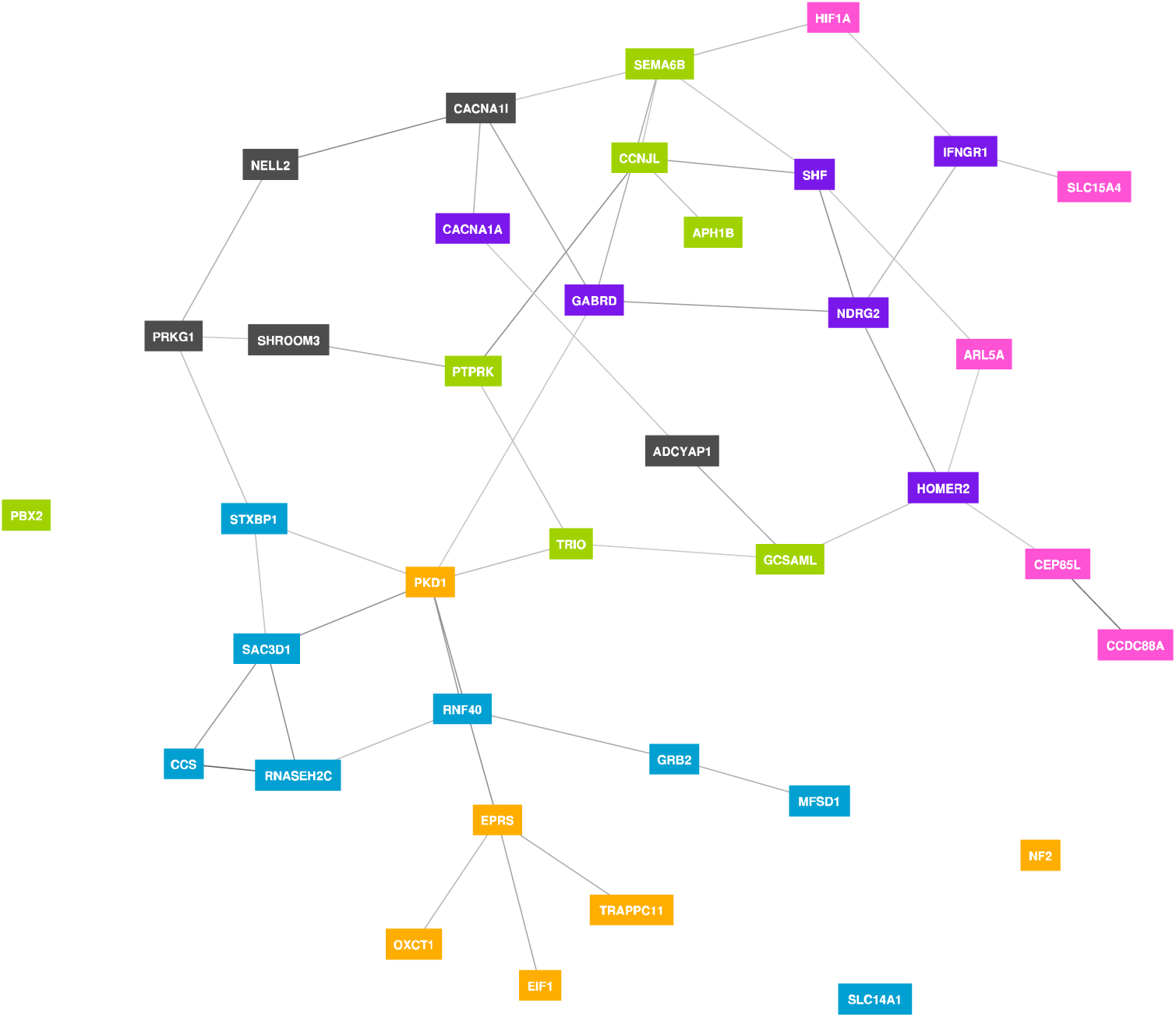
Gene clustering of differentially expressed genes for the cerebellar cortex based on gene co-expression. Public RNA sequencing data (N=31,499) was used to determine coexpression profiles. Gene cluster 1 identified in blue. Gene cluster 2 identified in green. Gene cluster 3 identified in purple. Gene cluster 4 identified in orange. Gene cluster 5 identified in pink. Gene cluster 6 identified in black.

### Expression levels across different tissue types and brain developmental stages

Expression levels in GTEx53 show that most of the DEGs are highly expressed in the brain cerebellar hemisphere and brain cerebellum (Supplementary Figure 3). Additionally, different regions of the brain have different expression levels for these genes. Expression levels across 29 different ages of brain samples from BrainSpan show that most of the DEGs are stably expressed during development (Supplementary Figure 4).

### Genome-wide gene association study of previous ET GWAS data identifies calcium-relevant pathways

To narrow down relevant pathways, a genome-wide gene association study (GWGAS) was done. The input SNPs were mapped to 18,220 protein coding genes. The genome-wide gene association study (GWGAS) identified *BUB1* reaching Bonferroni genome-wide significance and several genes reaching suggestive significance (Figure 3A). Clustering of the genes based on co-expression identified four distinct clusters (Supplementary Figure 5). From the pathway analyses, calcium ion-regulated exocytosis of neurotransmitters in GO was significantly enriched in the pathway in cluster three (P=9.7E-03). Gene-level enrichment analyses in GTEx 30 v7 found the brain to be significantly associated (P=0.001). After, the enrichment was queried in the brain-relevant regions of GTEx 53 v7 we found the cerebellar cortex and frontal cortex to pass Bonferroni-corrected significance (Figure 3B). No ET GWAS SNPs were found to be eQTLs for the dysregulated genes based on the GTEx53 database (Supplementary Figure 6-12).

**Figure 3.**
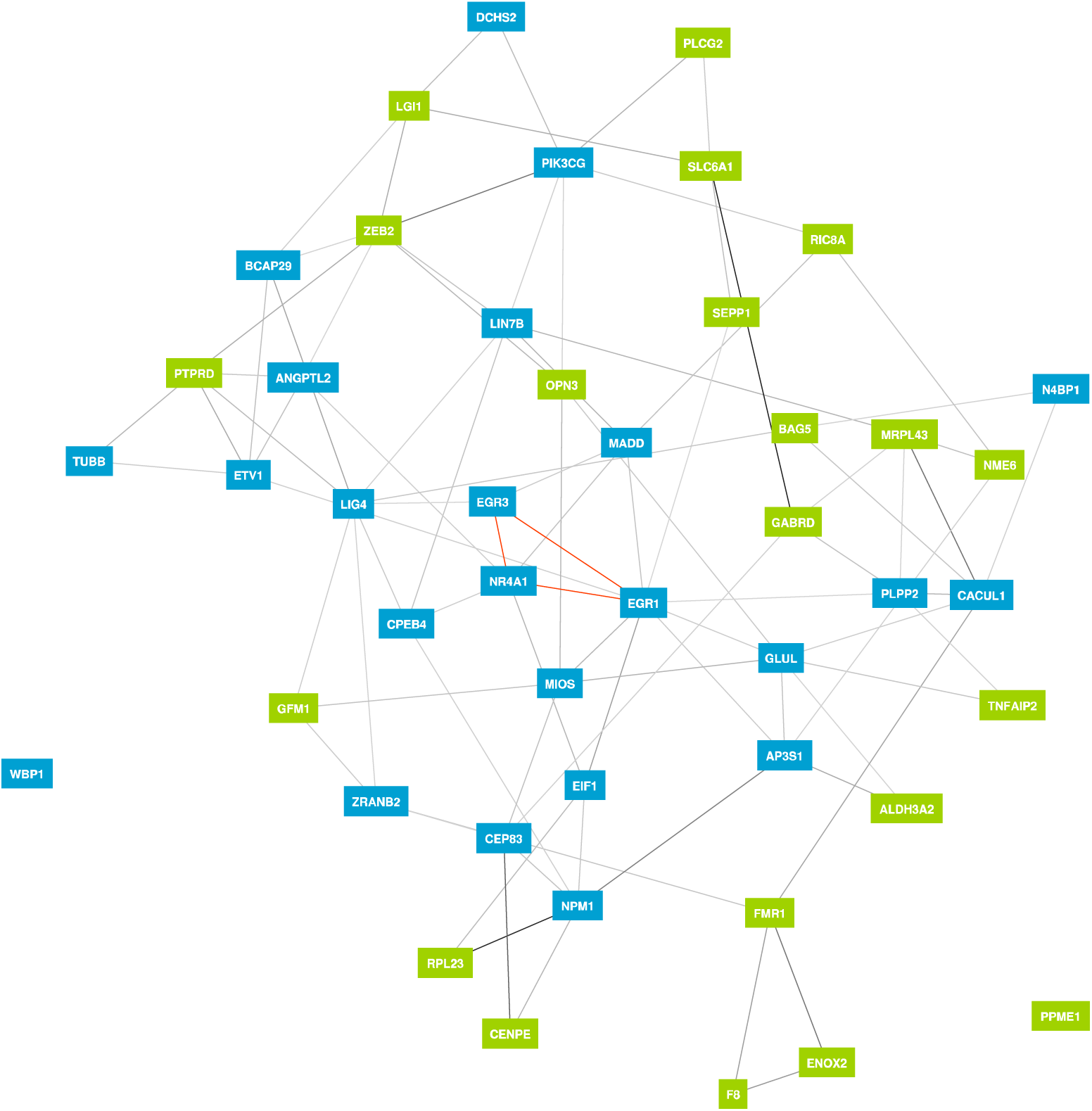
Gene clustering of differentially expressed genes for the dentate nucleus based on gene co-expression. Public RNA sequencing data (N=31,499) was used to determine co-expression profiles. Gene cluster 1 identified in blue. Gene cluster 2 identified in green.

**Figure 3.**
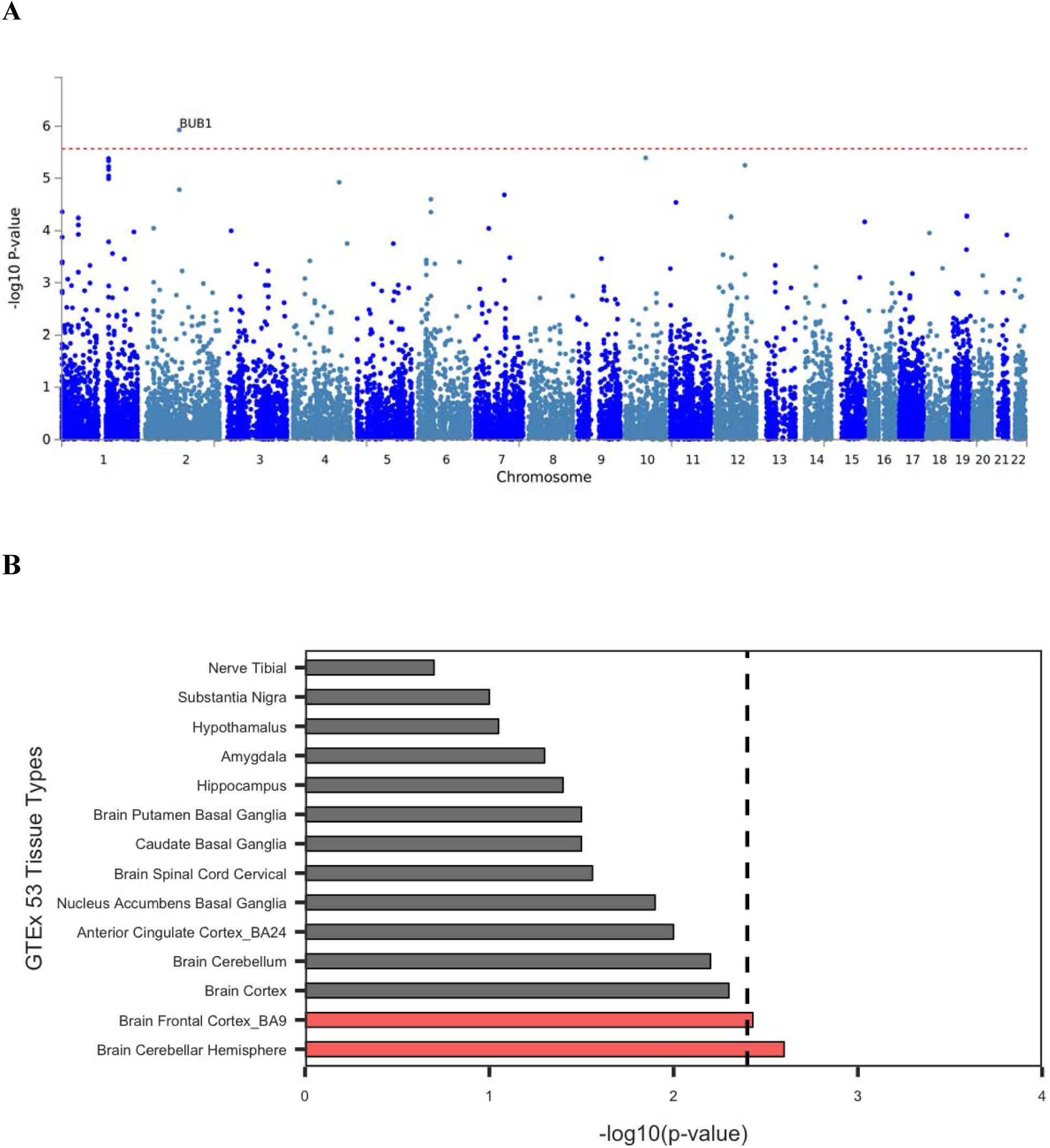
Genome-wide gene Manhattan plot and brain-relevant tissue enrichment profile. **(A)** Manhattan plot of gene-level associations (N=7,154). Bonferroni significance threshold at 2.744E-06 shown with the red dashed line in the plot. **(B)** Gene-level enrichment analysis of GWGAS genes in brain-relevant tissue of GTEx53. Bonferroni-corrected significance set at 0.0036, indicated by the dashed line.

### Phenome-wide association of DEGs

The phenome-wide association study (pheWAS) showed that *SHF* was significantly associated with blood pressure medication (P=9.30E-08) and body mass index (BMI) (P=1.5E-07) (Figure 4). No other DEGs had associations that passed a Bonferroni-corrected p-value.

**Figure 4.**
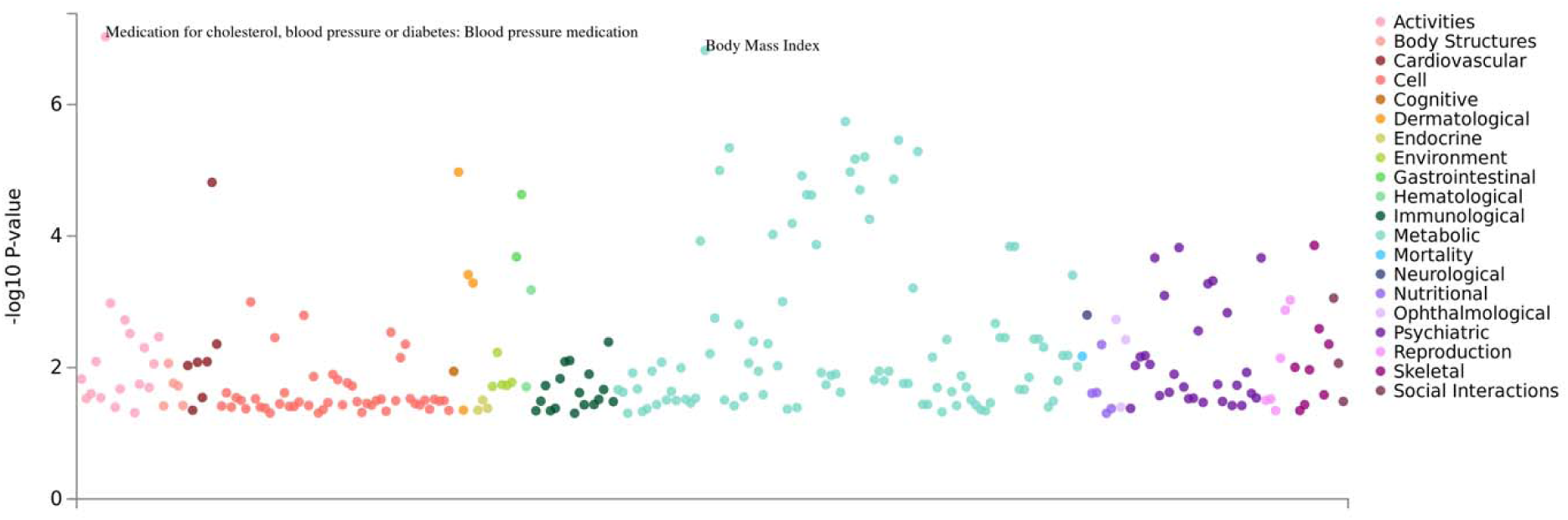
PheWAS manhattan plot of *SHF* for different domains. Colours indicate respective domains. Only p-values less than 0.05 were plotted.

## Discussion

Interestingly, calcium pathways and relevant genes were significant for differential expression and pathway analyses. In a paper by Topaktas *et al*. (1987), the use of calcium blockers led to intensified tremors in ET patients^7^. Interestingly, the calcium channel gene, *CACNA1A*, had lower levels of expression in ET patients, suggesting that cellular calcium may be relevant to the ET phenotype. In mice, knockout *CACNA1A* lines shown a tremor phenotype according to The Jackson Laboratory mice database. Furthermore, *CACNA1A* has been shown to be highly expressed in Purkinje cells, a relevant ET cell type^8^. The translated protein of *PRKG1*, PKG, has been reported to increase opening of calcium-activated potassium channels, further indicative of the relevance of calcium in ET^9^. Furthermore, irregular GABA-A receptor function has been previously shown to be affected by *CACNA1A*^10^. It is hypothesized that defective GABA receptors contribute towards the ET phenotype by disinhibition of cerebellar pacemaker output^11^. Additionally, the gene has been shown to be highly co-expressed with *GABRA4* based on the Gene Network database (P=1.6E-13), reinforcing *CACNA1A* as a gene of interest for ET.

The top DEG, *PRKG1*, has been shown to regulate cardiovascular and neuronal health^9^. Specifically, the RNA isoform two (PRKG1B) was most significantly differentially expressed at the transcript level. Currently, there has yet to be any study to link *PRKG1* and ET. However, the gene is highly expressed in Purkinje cells, which is a relevant cerebellar cell type in ET^12^. Also, the ET brain staining did not show significant Purkinje cell loss, suggesting transcriptomic dysregulation of Purkinje cells may be more relevant to ET than degeneration. Interestingly, *PRKG1* has been associated with alcohol misuse and many ET patients report reduced tremor intensity with alcohol, however, the pheWAS data did not show any associations^13^.

Pathway analyses additionally found several potentially relevant pathways such as axon guidance and neuromuscular junction. Interestingly, a possibly deleterious variant in *TENM4*, a regulator of axon guidance, was shown to segregate in ET families and cause tremor in knockout mice, reinforcing the relevance of this pathway in ET^14^.

The GWGAS identified *BUB1* as a significantly enriched gene for ET. *BUB1* is transcribed and translated into a serine/threonine kinase, similar to *STK32B*, which was a significant ET locus previously identified^15^. Amongst the significantly enriched pathways in the GWGAS and DEGs, calcium ion-regulated exocytosis of neurotransmitter was found to be in common between the two. This suggests that dysregulation in calcium homeostasis may affect relevant neurotransmitter exocytosis in ET patients.

The relationship of *SHF* with blood pressure medication from the pheWAS may suggest that beta-blockers interact with *SHF*. Beta-blockers such as propranolol can reduce kinetic tremor in certain ET patients^16^. The gene was clustered with the calcium-related genes such as *CACNA1A*, suggesting that beta-blockers may influence pathways identified from that cluster such as calcium channel activity. Furthermore, with the associated BMI trait, it could suggest that it is a potential risk factor for ET. Future studies should investigate the relationship between this gene and those phenotypes and determine whether *SHF* may be a biomarker for beta-blocker responsive ET patients.

Interestingly, examination of the dentate nucleus and cerebellar cortex from the same individuals revealed them to have distinct transcriptome profiles—the top DEGs were different between the two tissues. Based on the GTEx53 expression profiles of different tissue types, the differential expression across tissues reinforces the notion that the dentate nucleus and cerebellar cortex would have different DEGs. The BrainSpan database showed that the DEGs are similarly expressed across different ages and that adulthood has moderate to high expression of the DEGs.

Olfactory transduction and signaling were pathways enriched in both the dentate nucleus and cerebellar cortex. Past studies have had conflicting views on whether olfactory loss is an endophenotype for ET. However, the transcriptomic data objectively supports that a subset of ET patients likely have dysregulated olfactory phenotypes. Furthermore, in *MAPK* pathways were enriched in the dentate nucleus. This is interesting because beta-blockers have been shown to have downstream effects on MAPK-relevant pathways.

Here, we report the first transcriptomic study of ET and identified several dysregulated genes and relevant pathways. However, we acknowledge that bulk RNA sequencing cannot thoroughly distinguish which cell types may be driving the differentially expressed signals. Further replication studies investigating the transcriptome should be done for ET as the disease is highly heterogeneous. Future studies could investigate the spatial transcriptomics or single-cell sequencing of ET relevant tissue and integrate large datasets to further refine relevant genes and pathways.

## Methods

### Sample selection and criterion

Patients attending MDCS (Movement Disorder Clinic Saskatchewan) are offered autopsy at no cost to the family/estate. Autopsy consent is granted by the next-of-kin after death of the patient. The body is transported to Saskatoon and autopsy is performed within 24 h of death. The autopsy consent is approved by the Saskatoon Health Region Authority and the use of brain for research is approved by the Bioethics Board of the University of Saskatchewan. Further details on patient recruitment can be found from Rajput et al. (2015) and Rajput et al. (2016)^17,18^.

ET brains (N=16) were dissected to obtain the dentate nucleus and cerebellar cortex. Samples were selected based on the following criteria: grossly unremarkable cerebellum, staining showed no noticeable degeneration, no other neurological disorders or movement disorders (i.e. Parkinson’s disease or dystonia), no signs of dementia or mild/moderate Alzheimer’s changes, and have definite or probable ET. Controls (N=16) were age- and sex-matched. Additionally, the controls did not have any noticeable neurodegenerative or psychiatric disorders. The pH of all brains was neutral. Samples were matched to have an average RIN of 5.

### RNA extraction and sequencing

RNA was extracted from 64 samples using the RNeasy Lipid Tissue Mini Kit (Qiagen). Two samples were removed due to low quality. The RNA concentration was measured on the Synergy H4 microplate reader. RNA was sent to Macrogen Inc. for sequencing. Library preparation was done using the TruSeq Stranded Total RNA Kit (Ilumina) with Ribo-Zero depletion. Sequencing was done on the NovaSeq 6000 at 150bp paired-end reads with a total of 200M reads. Samples were randomized for tissue dissection, RNA extraction, Ribo-Zero depletion, library preparation and sequencing to account for potential batch effects.

### Data processing, differential expression, splicing and eQTL analyses

The FASTQ files were pseudo-aligned using Salmon using the Ensembl v94 annotation of the human genome^19^. For data processing and parameters of Salmon, please refer to Liao et al. (2019)^15^. Sleuth was used to identify DEGs^20^. The data was analyzed with the following full model for the likelihood ratio test: Gene expression ∼ disease status + sex + age + sex:disease status + age:sex + sex:disease status. The reduced model for the likelihood ratio test is: Gene expression ∼ sex + age + sex:age. A Wald test was used to get beta values, which is a bias estimator. Beta approximates the extent at which estimated counts is affected by the disease status rather than technical and biological variability. It can be used to estimate the magnitude and direction of fold change. P-values were corrected using the Benjamini-Hochberg procedure to account for false discovery rate (FDR). Q-value (p-values corrected for FDR) significance was set for <0.05. Splicing analyses were done with SUPPA2 and rMATs^21,22^.

### Validation with reverse transcriptase qPCR

To validate the significant DEGs, reverse transcriptase qPCR (RT-qPCR) was done. Following the manufacturer’s protocol, the SuperScript Velo cDNA Synthesis Kit (ThermoFisher Scientific) was used to convert 1 µg of RNA into cDNA. A standard curve was made for each TaqMan probe to determine PCR efficiency. The gene *POLR2A* (polymerase [RNA] II [DNA-directed] polypeptide) was used as an endogenous control.

### Pathway enrichment analyses in brain tissue

Gene clustering was done using the GeneNetwork v2.0 RNA sequencing database (N=31,499). Clusters were independently analyzed for different enriched pathways in databases such as Reactome and GO. FUMA was used to identify enrichment in BrainSpan and GTEx 53 v7^23^. Tissue specificity was tested in GTEx 53 v7. Briefly, DEG sets were pre-made and the input genes were tested against the DEG sets using a hypergeometric test. DEGs with a q-value <0.30 were included for analysis.

### Gene-level analyses of ET GWAS data

The raw genotyping data from the discovery cohort of the 2016 ET GWAS was obtained (N=7154). Data quality control included the following: Hardy-Weinberg equilibrium t-test P < 1E-4, minor allele frequency MAF > 0.05, genotype missingness < 2%, and sex discordance. Imputation was performed using EAGLE2 and PWBT, using the Sanger Imputation Service, and samples with INFO < 0.3 were removed. The data was analyzed using BOLT-LMM with 15 principle components^24^. A genome-wide gene association study (GWGAS) was done using MAGMA using the European UK Biobank as a reference LD panel. Genome-wide gene association studies considers the combined association effect of all SNPs in a gene to aggregate into a combined gene-level P-value. Genes with a suggestive Bonferroni-corrected p-value (P<0.10) were further queried in downstream analyses. Gene expression analysis was done using MAGMA for GTEx 53 v7 and BrainSpan. In the 30-general tissue GTEx v7 and then looked at expression of all brain-relevant regions in GTEx 53 v7. Gene clustering was done using the GeneNetwork v2.0 RNA sequencing database (N=31,499)^25^. Pathway enrichment and gene ontology analyses were also done for GWGAS data. Additionally, all ET-associated GWAS loci (P<1E-06) were queried in GTEx53 to determine if any were eQTLs for the differentially expressed genes.

## Phenome-wide association of differentially expressed genes

To understand which phenotypes may be associated with the differentially expressed genes, a pheWAS was done using GWASAtlas, which uses public genome-wide association study (GWAS) data^26^. Bonferroni-corrected of 1.68E-05 (0.05/# of unique traits) was used. At the time, there were 2977 traits.

