## Supplementary Figures for "Multi-omics integration of the phenome, transcriptome and genome highlights genes and pathways relevant to essential tremor"

**Supplementary Figure and Tables**

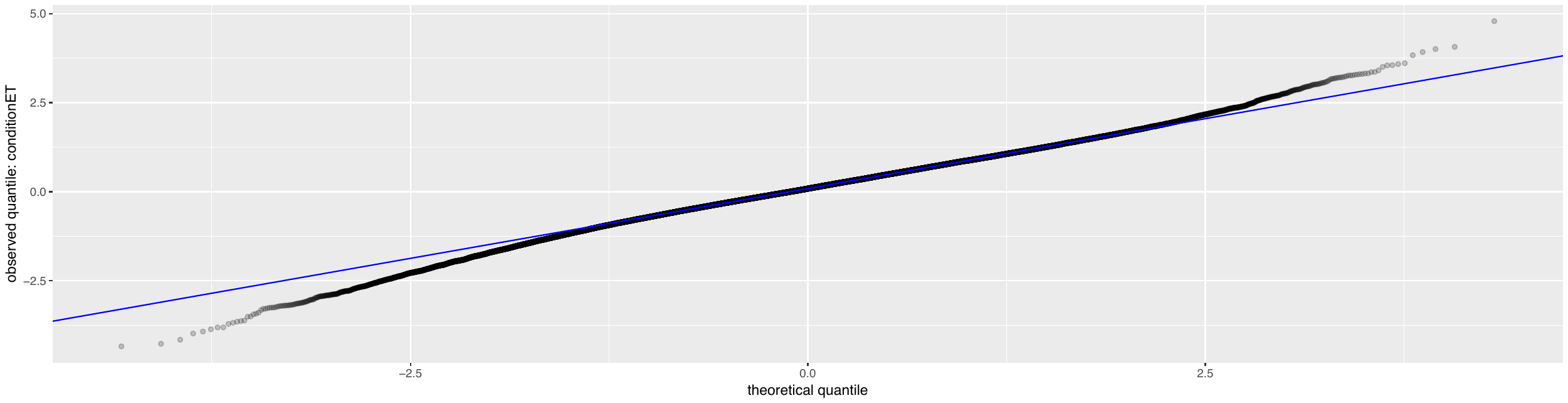

**Supplementary Figure 1. QQ-plot of cerebellar cortex differential expression data.** Blue line shows expected values.

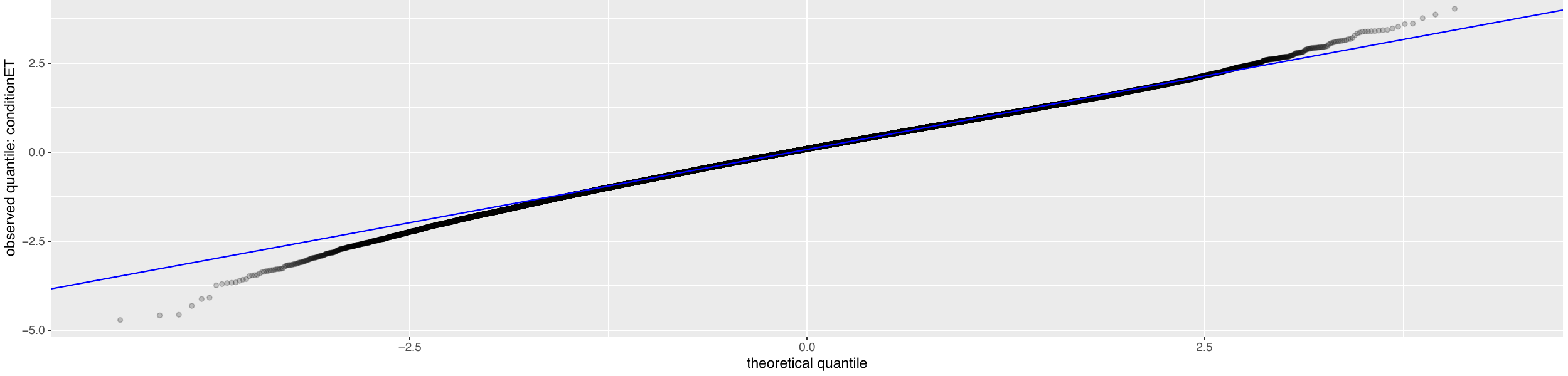

**Supplementary Figure 2. QQ-plot of dentate nucleus differential expression data.** Blue line shows expected values.

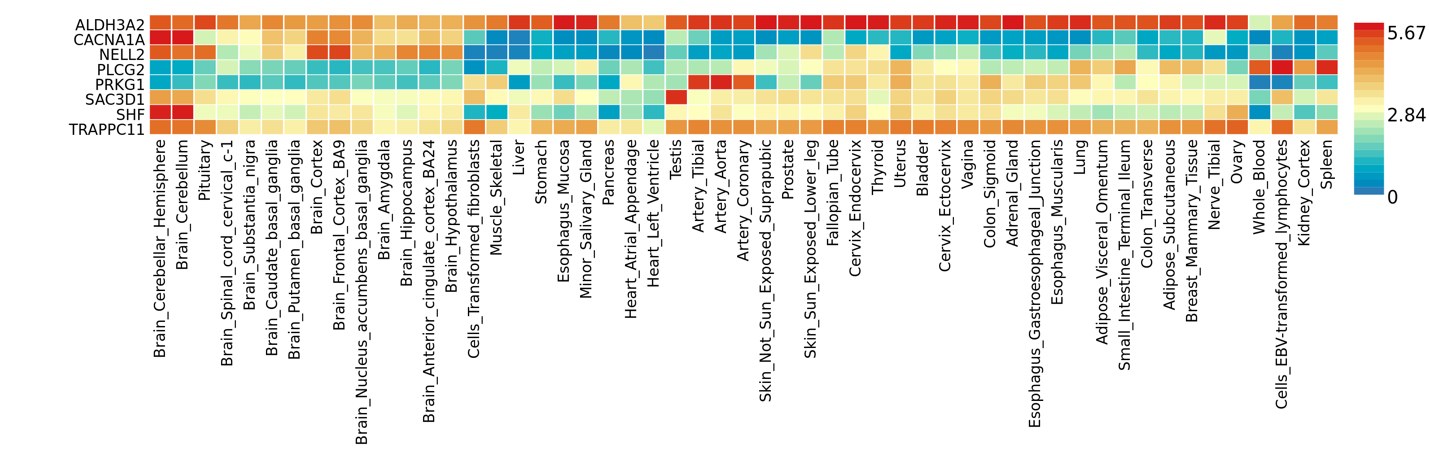

**Supplementary Figure 3. Expression of differentially expressed genes in GTEx53 tissue types.**

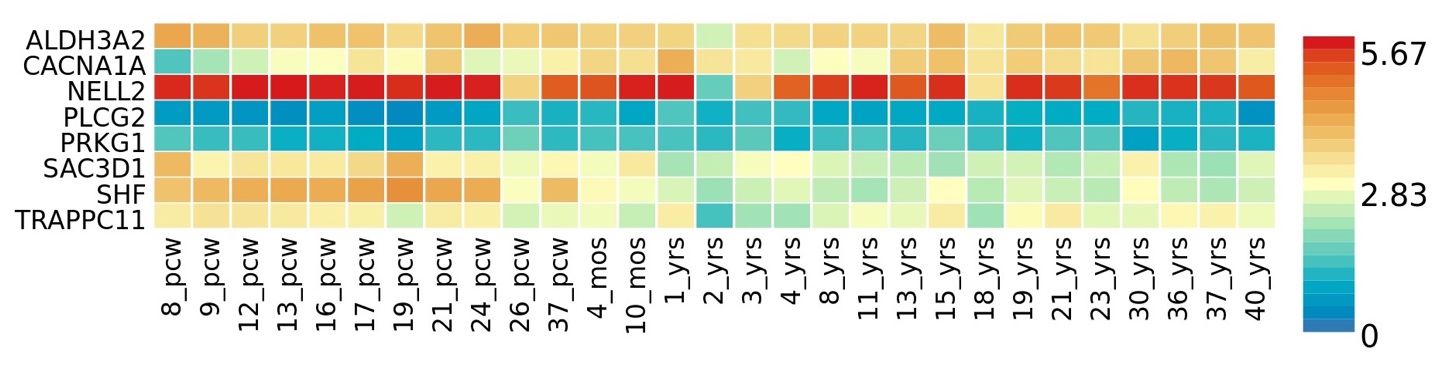

**Supplementary Figure 4. Expression of differentially expressed genes in BrainSpan 29 weeks of age dataset.**

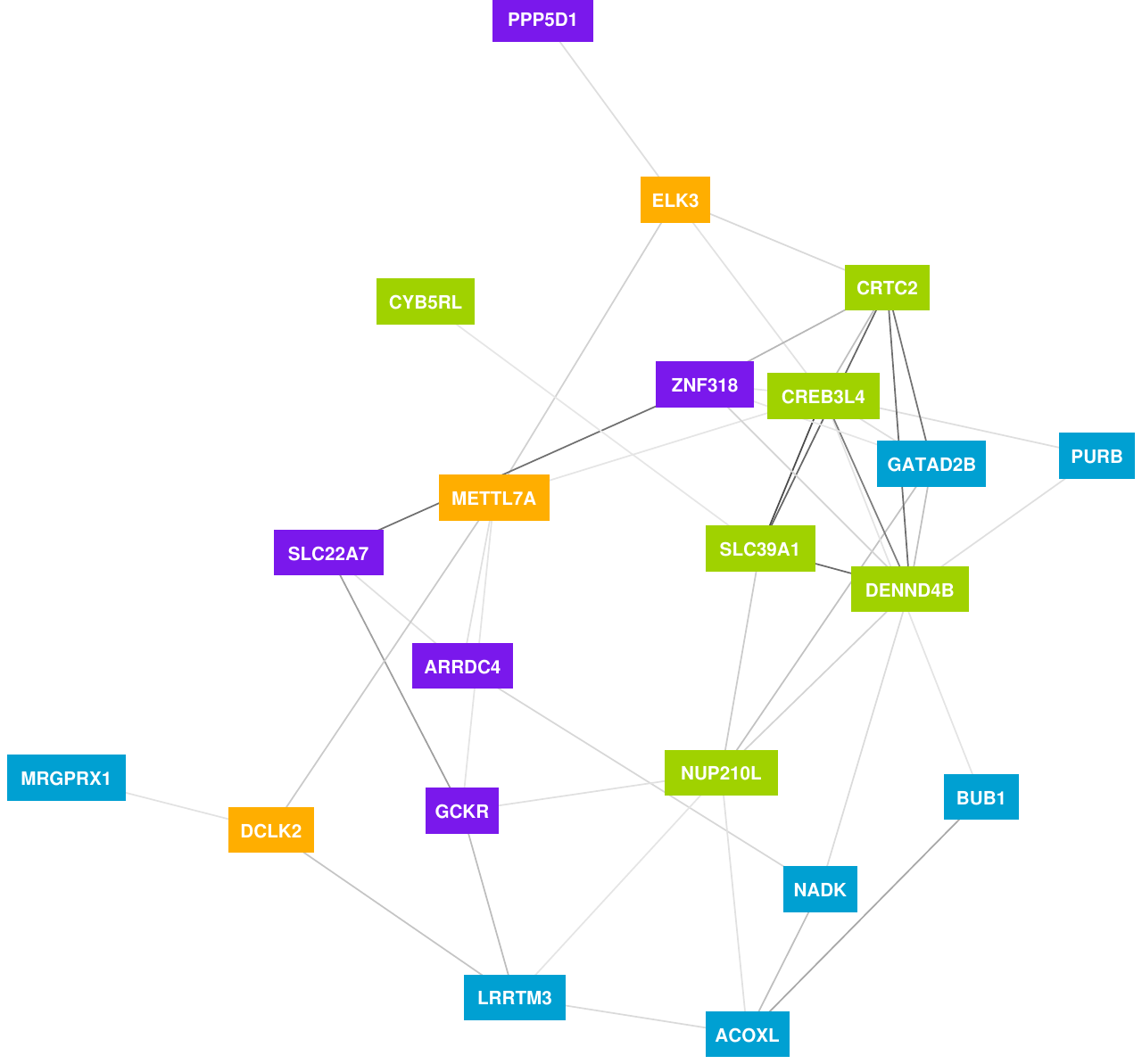

**Supplementary Figure 5. Gene clusters of GWGAS data.**

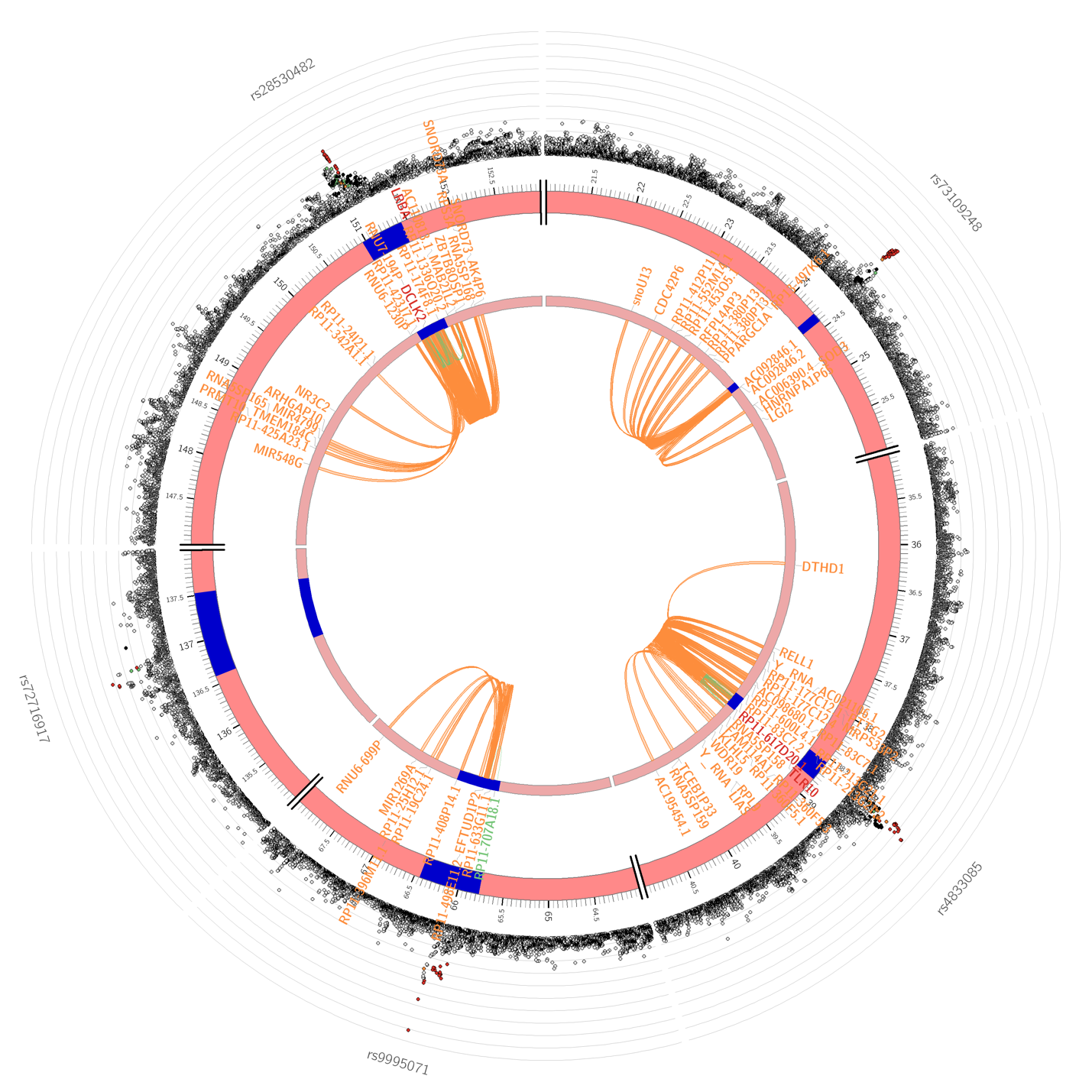

**Supplementary Figure 6. Circos plot of chromosome 4 for GWAS data showing eQTL and chromatin interaction of significant loci.**

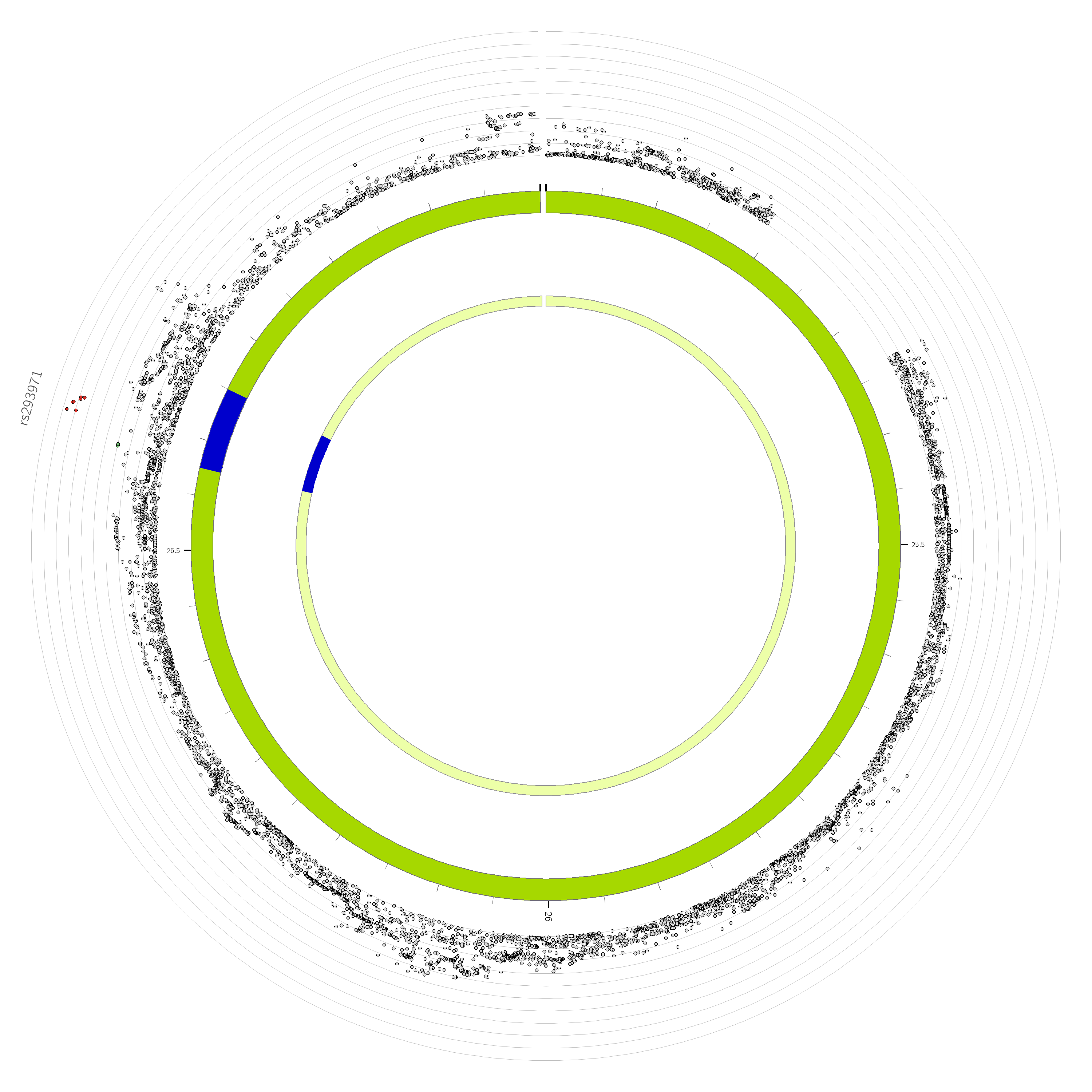

**Supplementary Figure 7. Circos plot of chromosome 11 for GWAS data showing eQTL and chromatin interaction of significant loci.**

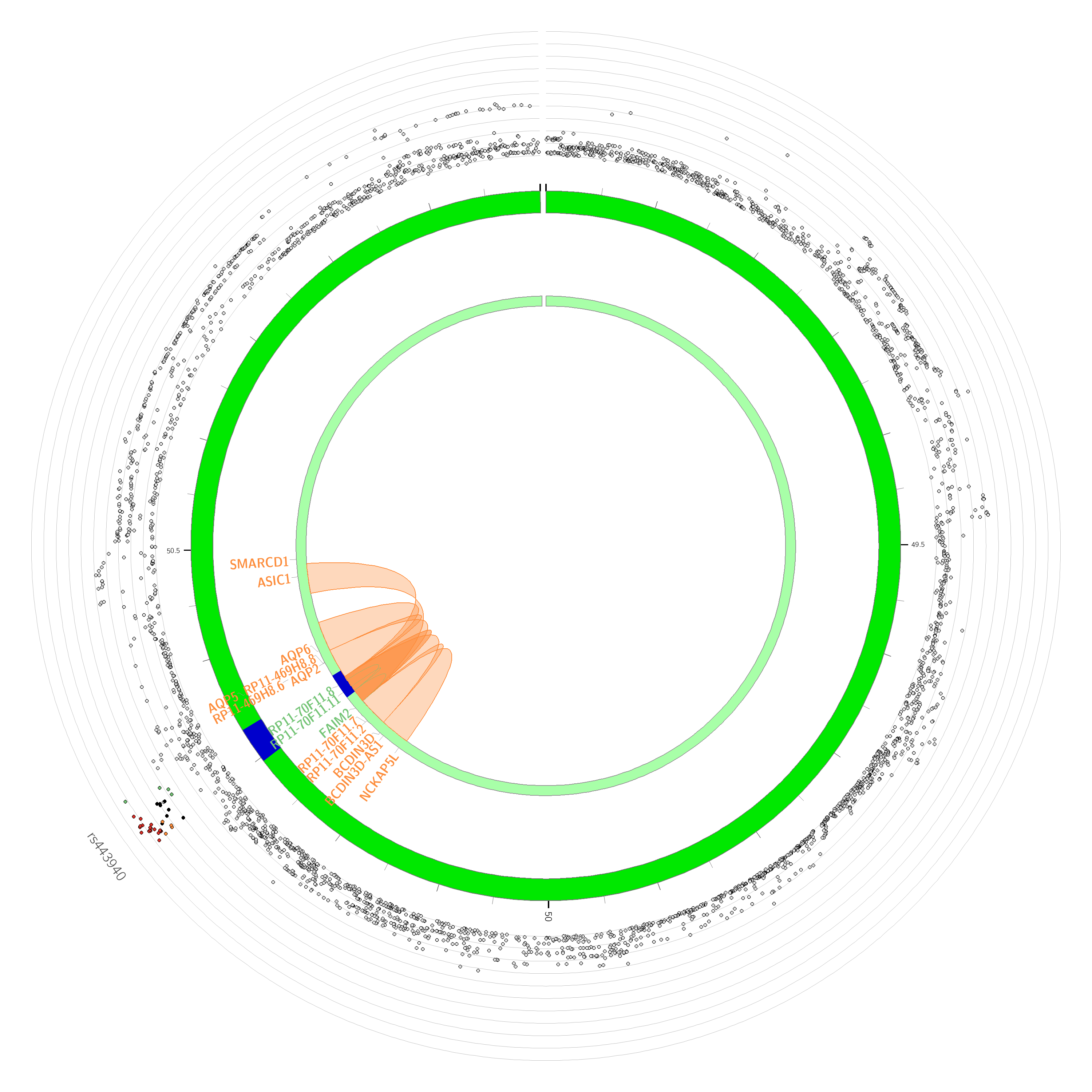

**Supplementary Figure 8. Circos plot of chromosome 12 for GWAS data showing eQTL and chromatin interaction of significant loci.**

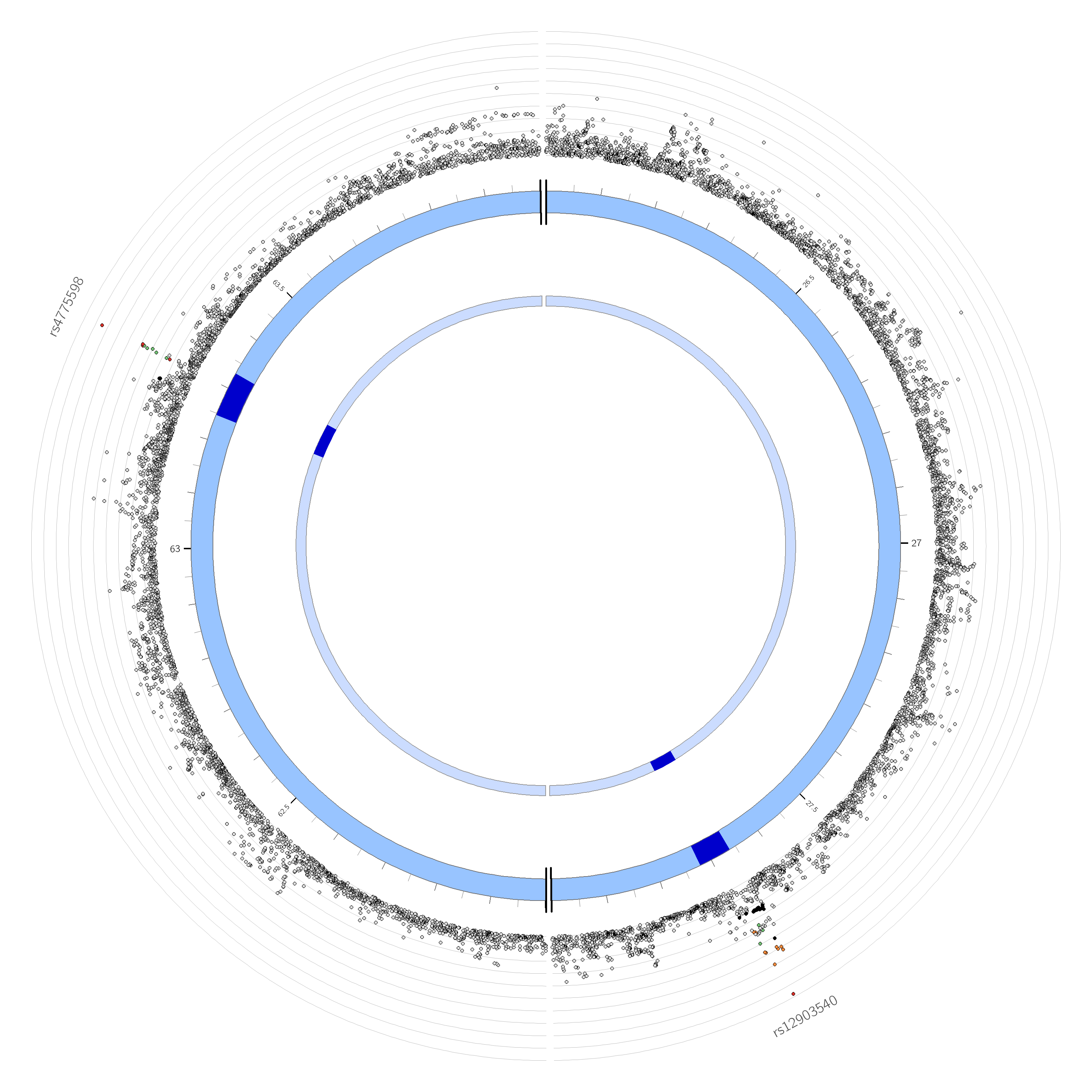

**Supplementary Figure 9. Circos plot of chromosome 15 for GWAS data showing eQTL and chromatin interaction of significant loci.**

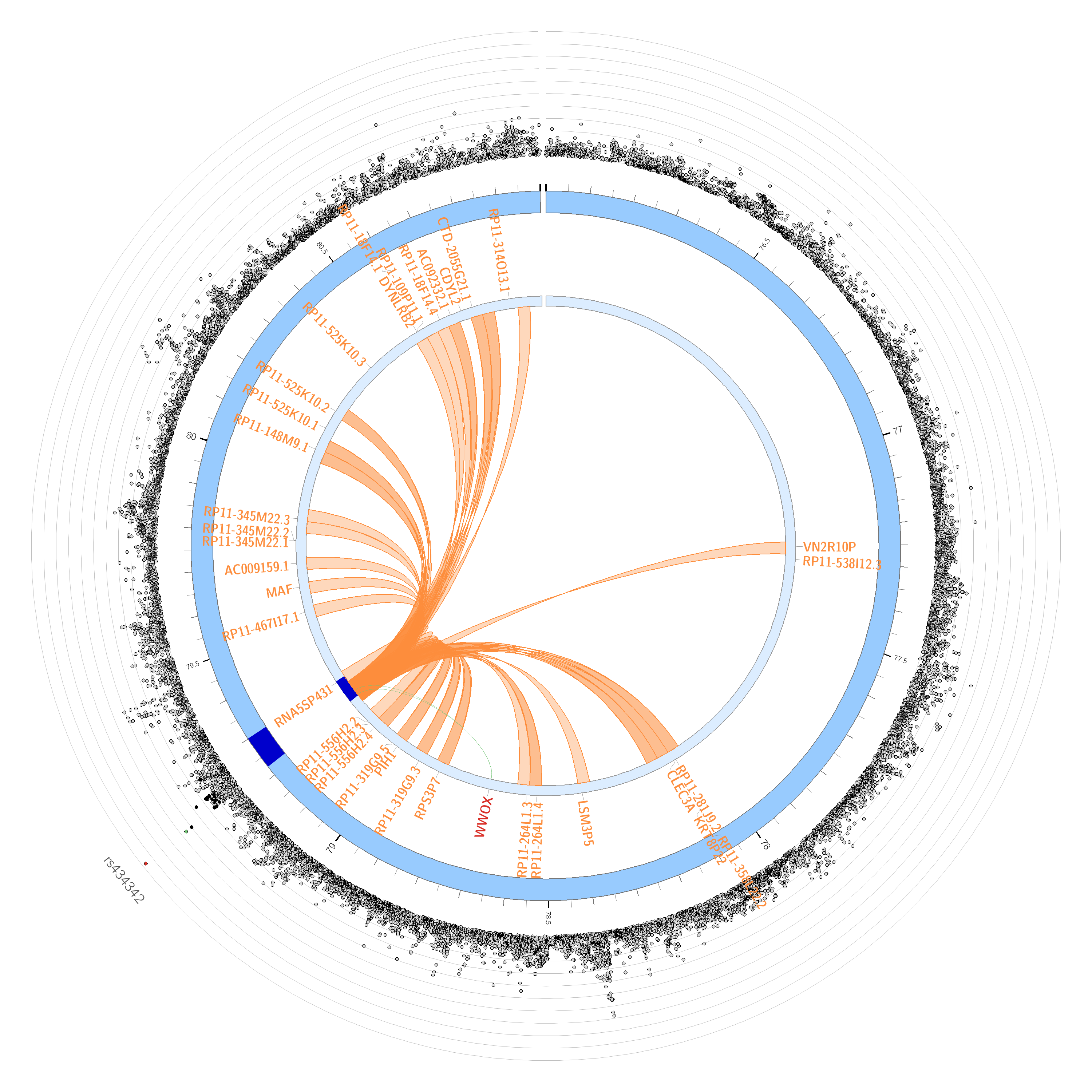

**Supplementary Figure 10. Circos plot of chromosome 16 for GWAS data showing eQTL and chromatin interaction of significant loci.**

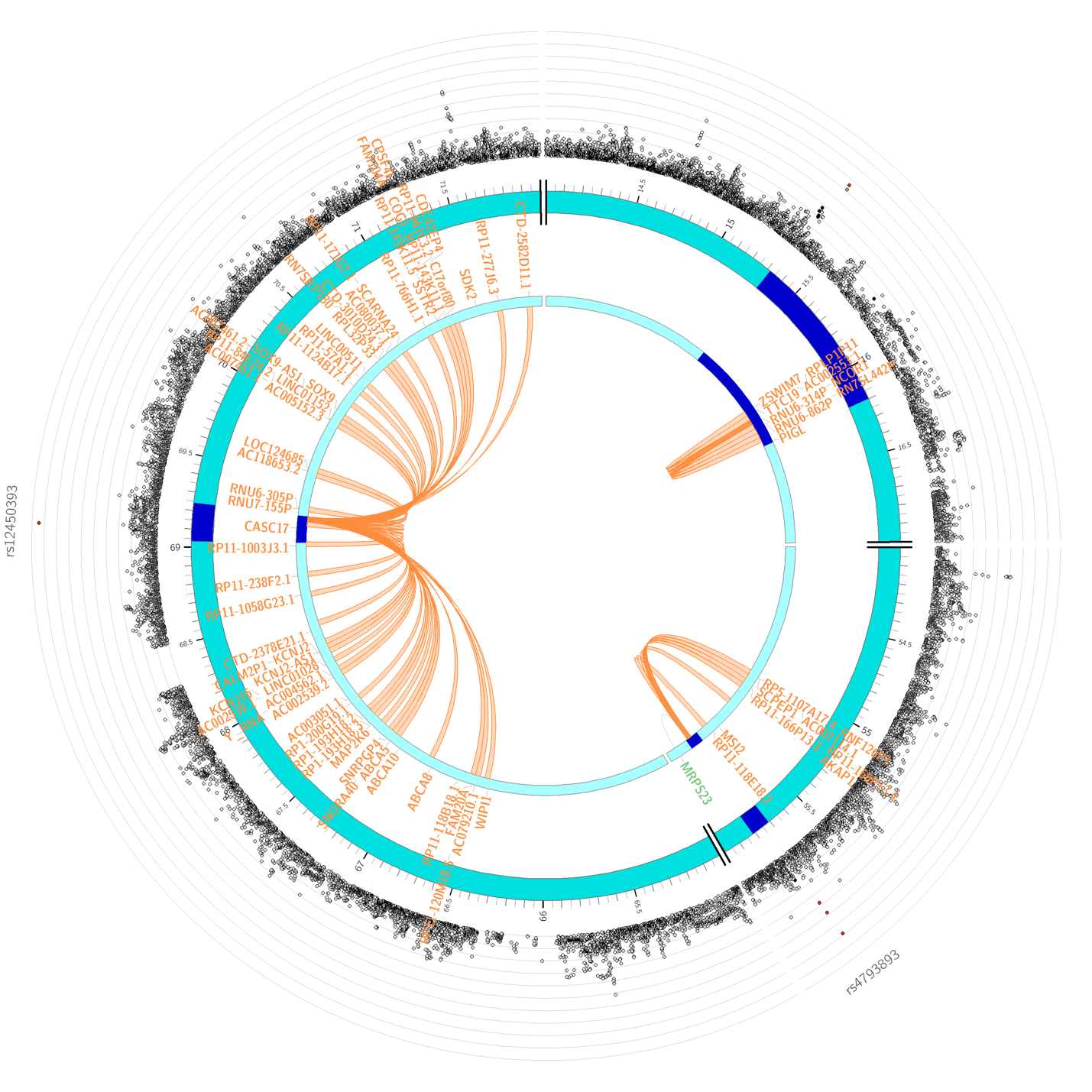

**Supplementary Figure 11. Circos plot of chromosome 17 for GWAS data showing eQTL and chromatin interaction of significant loci.**

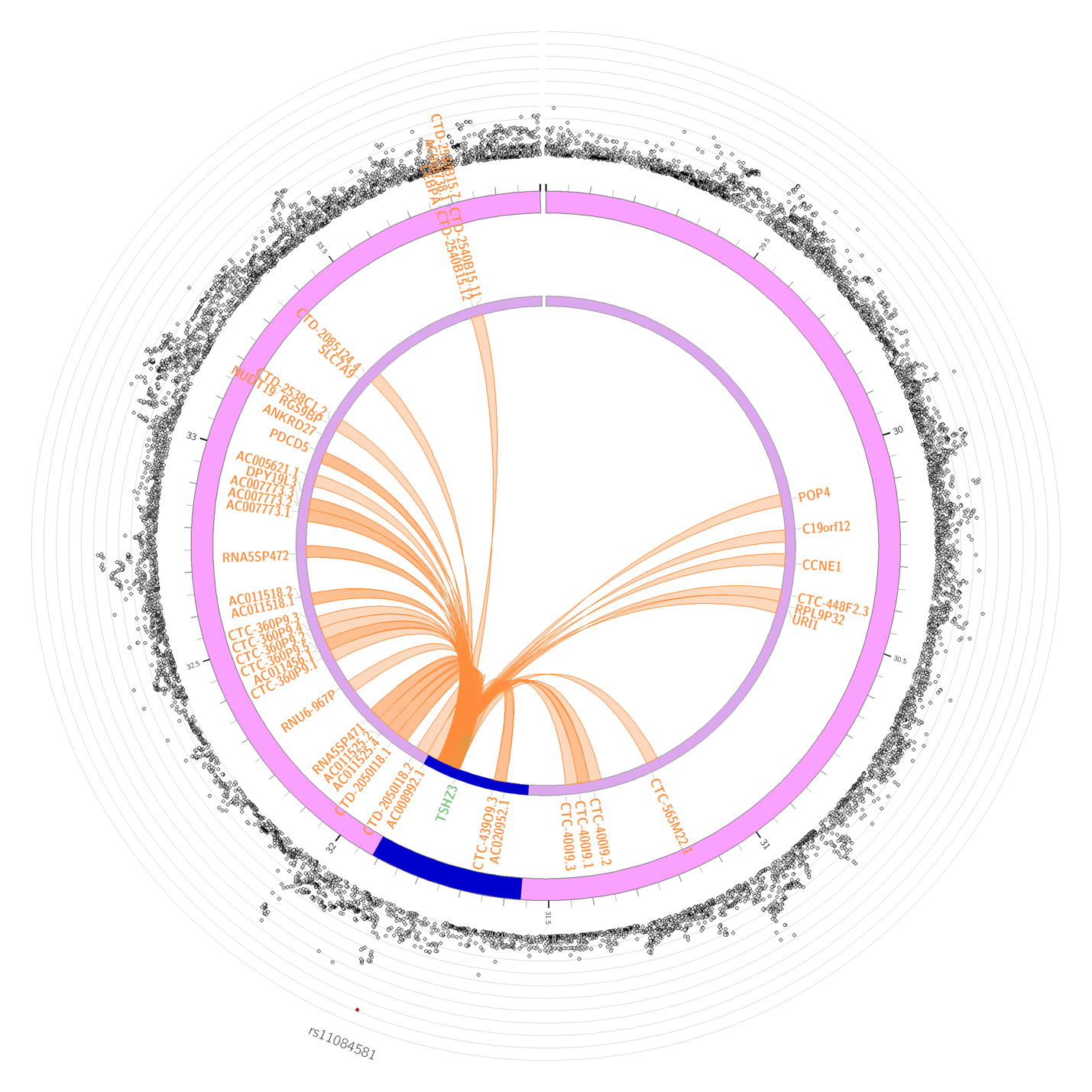

**Supplementary Figure 12. Circos plot of chromosome 19 for GWAS data showing eQTL and chromatin interaction of significant loci.**

**Supplementary Table 1.** Showing the sign concordance between RT-qPCR data and Wald statistic.

| **Gene** | **qPCR (∆∆ct)** | **RNAseq Wald statistic** |
| --- | --- | --- |
| *CACNA1A* | -0.113 | -15.30147 |
| *NELL2* | -0.868 | -8.601377 |
| *PRKG1* | -0.77 | -11.89647 |
| *ALDH* | 0.22 | 21.8715 |
| *PRKG1* | -0.424 | -11.89647 |
| *TRAPPC11* | -0.223 | -16.59086 |
| *PLCG2* | -0.389 | -9.73396 |
| *SHF* | -0.135 | -17.69 |
